# Fast EEG-based decoding of the directional focus of auditory attention using common spatial patterns

**DOI:** 10.1101/2020.06.16.154450

**Authors:** Simon Geirnaert, Tom Francart, Alexander Bertrand

## Abstract

**Objective:** Noise reduction algorithms in current hearing devices lack information about the sound source a user attends to when multiple sources are present. To resolve this issue, they can be complemented with auditory attention decoding (AAD) algorithms, which decode the attention using electroencephalography (EEG) sensors. State-of-the-art AAD algorithms employ a stimulus reconstruction approach, in which the envelope of the attended source is reconstructed from the EEG and correlated with the envelopes of the individual sources. This approach, however, performs poorly on short signal segments, while longer segments yield impractically long detection delays when the user switches attention.

**Methods:** We propose decoding the directional focus of attention using filterbank common spatial pattern filters (FB-CSP) as an alternative AAD paradigm, which does not require access to the clean source envelopes.

**Results:** The proposed FB-CSP approach outperforms both the stimulus reconstruction approach on short signal segments, as well as a convolutional neural network approach on the same task. We achieve a high accuracy (80% for 1 s windows and 70% for quasi-instantaneous decisions), which is sufficient to reach minimal expected switch durations below 4 s. We also demonstrate that the decoder can adapt to unlabeled data from an unseen subject and works with only a subset of EEG channels located around the ear to emulate a wearable EEG setup.

**Conclusion:** The proposed FB-CSP method provides fast and accurate decoding of the directional focus of auditory attention.

**Significance:** The high accuracy on very short data segments is a major step forward towards practical neuro-steered hearing devices.

## I. Introduction

Current hearing aids, cochlear implants, and other assistive listening devices contain noise reduction algorithms to assist people suffering from hearing deficits. However, these algorithms often fail in so-called ‘cocktail party’ scenarios where multiple speakers or other sound sources are simultaneously active. This is not only because the noise suppression becomes more difficult, but primarily because the hearing device lacks information about which source the user *intends* to attend and which other sources need to be treated as background noise.

Instead of using unreliable heuristics, such as selecting the most frontal source or the source with the highest intensity, one could try to extract the attention-related information directly from where it originates, i.e., the brain. This is known as auditory attention decoding (AAD). The development of AAD algorithms that process brain signals to, e.g., steer a speech enhancement algorithm towards the attended speaker in a mixture of other speakers, could lead to a new class of so-called ‘neuro-steered’ hearing devices [1], [2], improving the quality of life of people suffering from hearing deficits.

The discovery that the cortical activity follows the envelope of the attended speech stream [3]–[5] is in this context crucial. This insight laid the foundation of a first class of AAD algorithms based on non-invasive neural recordings from magneto- or electroencephalography (MEG/EEG). These algorithms typically employ a stimulus reconstruction approach in which a decoder reconstructs the attended speech envelope from the EEG. The decoded envelope is then correlated with the speech envelopes of the individual speakers. The speaker corresponding to the highest correlation coefficient is identified as the attended speaker. This algorithm was proposed for the first time in [6], using a linear minimal-mean-squared-error-based decoder. Later, other variations of this AAD algorithm were developed, of which an overview can be found in [2].

AAD algorithms using the stimulus reconstruction approach, however, all suffer from the same limitations:

1. The stimulus reconstruction approach takes too long to make a reliable decision. The AAD accuracy (the percentage of correct decisions) drastically decreases with shorter decision windows, especially below 10 s [6]–[8]. A decision window corresponds to the signal length over which the correlation coefficients between the EEG-decoded envelope and the original speech envelopes are estimated, where short decision windows result in unreliable correlation estimates. This results in a speed-accuracy trade-off. In [8], it is shown that short decision window lengths are favorable in the context of robust AAD-based gain control during dynamic switching, even if they have a lower accuracy. Nevertheless, due to the low accuracy for these short decision window lengths, it theoretically takes more than 15 s to establish a reliable and controlled gain switch to the new attended speaker after the user switches attention [2], which is impractically long for neuro-steered hearing device applications. The stimulus reconstruction approach inherently suffers from this limited performance due to the decoding of a low-frequency envelope, which contains relatively little information per second, as well as due to the low signal-to-noise ratio of the neural response to the stimulus in the EEG.
2. The stimulus reconstruction approach requires the (clean) individual speech envelopes. Although several attempts have been made to combine speech separation algorithms with AAD [1], [9]–[11], the demixing of all speech envelopes adds a lot of overhead, and the demixing process often negatively affects AAD performance or may even completely fail in practical situations.

In this paper, we employ a new paradigm that avoids these limitations, focusing on decoding the directional focus of attention from the EEG, rather than directly identifying the attended speaker. Inherently, this avoids the need of demixing the speech mixtures into its individual contributions. Moreover, we hypothesize that this paradigm will improve AAD accuracy for short decision window lengths, as it is based on brain lateralization, which is an instantaneous spatial feature, rather than a correlation-based temporal feature.

This new AAD paradigm is justified by recent research that shows that the auditory attentional direction is spatio-temporally encoded in the neural activity [4], [12]–[18], ergo, that it could be possible to decode the spatial focus of attention from the EEG. In [19], an AAD algorithm based on a convolutional neural network (CNN) has been established to decode the spatial locus of attention in a competing speaker scenario, which showed very good results on short decision windows (76.1% accuracy on 1 s decision windows). However, this CNN-based approach shows high inter-subject variability and requires large amounts of training data (e.g., data of other subjects in combination with subject-specific data as in [19]) in order to train a subject-specific decoder. Therefore, in this paper, we focus on data-driven *linear* filtering techniques, which typically require less training data, are more robust and stable, and are computationally cheaper, as well as easier to update. More specifically, we exploit the direction-dependent spatio-temporal signatures of the EEG using (filterbank) common spatial pattern (FB-CSP) filters, which are popular in various brain-computer interface (BCI) applications [20], [21].

In Section II, we concisely introduce the (FB-)CSP classification pipeline to determine the directional focus of attention. In Section III, we describe the data used to run experiments, the concrete choices for the FB-CSP filter design, and the performance metrics to transparently and statistically validate the experiments that are reported and analyzed in Section IV. Conclusions are drawn in Section V.

## II. Decoding direction of attention using CSPs

In this section, we review the CSP procedure [20] to decode the directional focus of attention. CSP filtering is one of the most popular techniques used for spatial feature extraction in BCI applications, e.g., in motor imagery [20]–[22]. The goal is to project multichannel EEG data into a lower-dimensional subspace that optimally discriminates between two conditions or classes. This is established by optimizing a spatial filter in a data-driven fashion, which linearly combines the different EEG channels into a few signals in which this discriminative property is maximally present.

For the sake of an easy exposition, we first define CSP filtering for a binary AAD problem, i.e., decoding whether a subject attends to one of two speaker positions, in Section II-A and II-B. In Section II-C, we explain how this can be generalized to more than two classes/directions. Finally, in Section II-D, we explain how the method can be applied to EEG data from unseen subjects without the need for any ground-truth labels on their auditory attention.

### A. CSP filtering

Consider a zero-mean *C*-channel EEG signal **x**(*t*) ∈ ℝ^*C×*1^, which can on each time instance *t* be classified into one of two classes 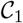 and 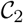 (e.g., attending the left or right speaker). The goal is to design a set of *K* spatial filters **W** ∈ ℝ^*C×K*^ that generate a *K*-channel output signal with uncorrelated channels *y*(*t*) = **W**^T^**x**(*t*) ∈ ℝ^*K×*1^, where the 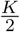 first filters maximize the output energy when 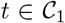, while minimizing the output energy when 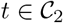, and vice versa for the other 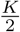 filters.

For example, the first column **w**_1_ of **W** results in 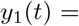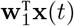, which should have a maximal output energy when 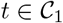 and a minimal output energy when 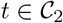:

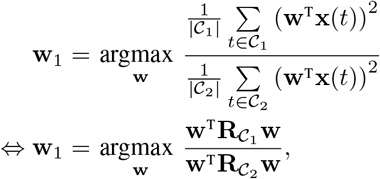

with 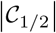 the number of time instances in 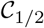 and

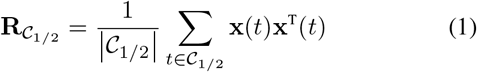

the sample covariance matrices of class 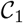 and 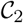. Fixating the output energy when 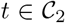, i.e., 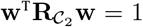, which is possible because **w** is defined up to a scaling, and solving the optimization problem using the method of Lagrange multipliers leads to the following necessary condition for optimality:

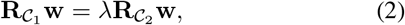

which corresponds to a generalized eigenvalue problem. It can easily be seen that the maximum is obtained for the generalized eigenvector corresponding to the largest generalized eigenvalue. A similar reasoning can be followed for **w**_*K*_, which maximizes, resp. minimizes the output energy when 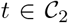, resp. 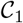, and is equal to the generalized eigenvector corresponding to the smallest generalized eigenvalue in (2). The other spatial filters can be found as the subsequent largest and smallest generalized eigenvectors. In its core essence, designing CSP filters thus corresponds to a joint diagonalization of the class-dependent covariance matrices [20].

### B. Classification using CSP filters

The CSP filtering technique can now be employed in a classification pipeline, in which a newly recorded EEG signal **x**(*t*) ∈ ℝ^*C×*1^, containing *C* channels, is classified into one of two classes, representing different directions of auditory attention (Fig. 1). The following sections describe the different components of this classification pipeline.

**Fig. 1:**
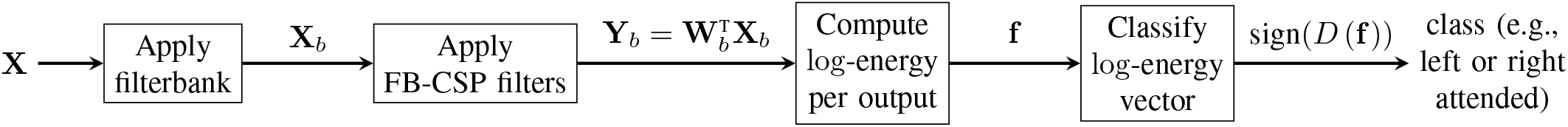
The FB-CSP filter outputs are used to generate the features that can be used to classify the EEG segment **X**.

#### 1) Filterbank CSP (FB-CSP)

Paramount for a well-performing CSP filtering is the selection of the appropriate frequency band related to the feature at hand. For the case of auditory attention decoding, one possibility is filtering in the *α*-band [4], [13], [14], [17], [18]. We here, however, do not want to make an a priori choice of the relevant frequency band(s). We thus adopt the so-called *filterbank* CSP (FB-CSP) technique, in which the EEG is first filtered into different frequency bands, after which the CSP filters are trained and applied per frequency band [20]–[22]. The filterbank thus results in *B* (number of frequency bands) filtered signals **x**_*b*_(*t*) ∈ ℝ^*C×*1^, one per frequency band *b* ∈ {*1*, …, *B*}, for all *C* EEG channels. The application of the pre-trained CSP filters per frequency band **W**_*b*_ ∈ ℝ^*C×K*^ results in *B K*-dimensional output signals 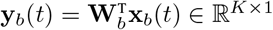.

An alternative extension, which is not pursued here, is the so-called common spatio-spectral pattern filter, in which the relevant frequency bands are determined fully data-driven, as a spatio-temporal filter is optimized to be maximally discriminative [23]. This comes, however, at the cost of an increase in parameters and corresponding problems with overfitting, in particular for high-density EEG data as used in this paper. These problems can partly be overcome by using more advanced regularization or dimensionality reduction techniques on the extended spatio-temporal covariance matrices (e.g., principal component analysis [24] or the pre-selection of relevant time lags to introduce sparsity). Furthermore, a different filter basis than the Dirac basis could be chosen to reduce the number of parameters or to incorporate expert knowledge [24].

#### 2) Feature extraction

The outputs of the FB-CSP filtering are now per decision window transformed into a feature vector **f** ∈ ℝ^*KB×*1^ that can be used for classification. This is typically done by computing the log-energy over these output signals per decision window [20], using a pre-defined decision window length *T*:

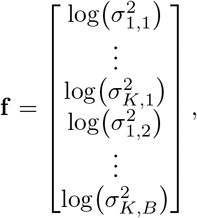

with the output energy 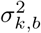 of the *k*^th^ output *y*_*k,b*_(*t*), for the *b*^*th*^ frequency band:

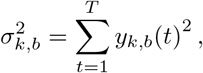

where *T* is the number of time samples in the decision window. Note that the decision window length *T* determines how much EEG data is used to make a single decision about the auditory attention of the subject. In a practical system, this will define the inherent delay to detect switches in attention.

#### 3) Classification

The feature vector **f** is used as the input for a binary classifier to determine the directional focus of attention. We here adopt Fisher’s linear discriminant analysis (LDA), which is traditionally used in combination with CSP filters [20]. In LDA, similarly to CSPs, a linear filter **v** ∈ ℝ^*KB×*1^ is optimized to provide the most informative projection. In this case, the most informative projection corresponds to maximizing the in-between class scatter, while minimizing the within-class scatter. This again leads to a generalized eigenvalue problem, which can, in this case, be solved analytically, leading to the following solution [25]:

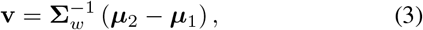

with **Σ**_*w*_ the covariance matrix of the features **f** computed across both classes, and *μ*_1/2_ the class (feature) means. Choosing the threshold or bias as the mean of the LDA projected class means leads to the following decision function:

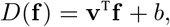

with **v** defined in (3) and bias

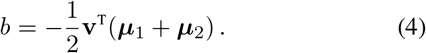

Finally, **f** is classified into class 1 if *D*(**f**) *>* 0 and into class 2 if *D*(**f**) *<* 0.

### C. Multiclass CSP classification

The classification scheme in Fig. 1 can be easily extended to a multiclass scenario, in which multiple directions of auditory attention are combined. This can be achieved by applying the strategy of Section II-A and II-B in combination with an appropriate coding scheme (e.g., one-vs-one, one-vs-all), both for the CSP and LDA step, or, by approximating a joint diagonalization of all the class covariance matrices at once in the CSP block [20], and only applying a coding scheme to the LDA step. Note that also the stimulus reconstruction approach is applicable for various directions/speakers [11], [26].

In this paper, we adopt the popular one-vs-all approach in BCI research [20]. In this approach, an FB-CSP filter and LDA classifier is trained for each direction to discriminate that particular direction from all the other directions. Given *M* directions (classes), this means that in (2), the 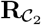 is replaced by 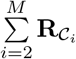, i.e., the sum of the covariance matrices of all classes except class 1. Correspondingly, an LDA classifier is trained to discriminate direction 1 from all the other directions. This is done for every other direction *m* ∈ {1, …, *M*}. Given *M* directions (classes), this thus results in *M* different CSP/LDA pairs.

In the end, for a new segment, the posterior probability of each classifier is computed using the multivariate normal distribution for the likelihood (which is assumed by LDA) and a uniform prior. To finally determine the correct class, the maximal posterior probability is taken over the *M* LDA classifiers.

### D. CSP classification on an unseen subject

The FB-CSP filters and LDA classifiers can be trained *subject-specifically*, meaning that the training is based on EEG data from the actual subject under test. However, in a neuro-steered hearing device application, this would require a cumbersome per-user calibration session where the subject is asked to attend to specific speakers with the intention to collect ground-truth labels to inform the FB-CSP filter design. To eliminate this requirement, one could train an AAD model in a *subject-independent* manner, meaning that data from subjects other than the test subject are used in the training phase, as done in [6] and [19] for the stimulus reconstruction and CNN approaches. This pre-trained model could then be ‘pre-installed’ on every neuro-steered hearing device, using it in a ‘plug-and-play’ fashion.

However, it is known from the BCI literature that the FB-CSP method often fails in such subject-independent settings due to too large differences in the spatial/spectral EEG patterns across different subjects [27]. To improve performance, the data from the subject under test can be used to modify the pre-trained subject-independent FB-CSP filters/LDA classifier. We adopt here two popular approaches to perform such adaptations, without requiring ground-truth labels for the data of the unseen test subject:

1. A very effective way of unsupervised updating of an LDA classifier for BCIs has been proposed in [28]. They conclude that simply updating the bias of the LDA classifier in (4) results in a significant improvement. Here, we update the bias of the subject-independently trained LDA with the unlabeled subject-specific features (resulting from the subject-independent FB-CSP filters), as this only requires the global mean, which is label-independent:

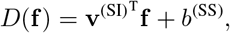

with subject-independent coefficients **v**^(SI)^ as computed in (3) on the data from all other subjects, and the subject-specific bias computed as:

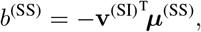

using the global mean *μ*^(SS)^ over all features **f**^(SS)^ of the new subject. The only requirement is that the subject-specific data on which the bias is updated is approximately balanced over both classes.
2. In [27], the authors found that a subject-independent FB-CSP method often fails, potentially because of the too high spectral subject-to-subject variability when using many narrow frequency bands. To overcome this issue, we replace the filterbank setting with a single filter (*B* = 1) to extract and pool a broader frequency range, of which the boundaries will be determined experimentally. This basically reduces the FB-CSP method to a CSP classification method with a prior bandpass filtering of the data. Note that only for the subject-independent experiments (Section IV-G), the FB-CSP method is reduced to a single frequency band. In all other subject-specific experiments, the FB-CSP approach is used.

## III. Experiments and evaluation

### A. AAD datasets

We apply the proposed FB-CSP classification method on two different datasets. The first dataset (Dataset I) has already been used extensively in previous work, mostly in the context of the stimulus reconstruction approach [2], [7], [8], [19], [29], and is publicly available [30]. It consists of 72 minutes of EEG recordings for each of the 16 normal-hearing subjects, who were instructed to attend to one of two competing (male) speakers. The competing speakers were located at −90° and +90° along the azimuth direction and there was no background noise. More details can be found in [7]. This dataset is used in all experiments, except those of Section IV-C and IV-D^1^.

The second dataset (Dataset II) consists of 138 minutes of EEG recordings for each of the 18 normal-hearing subjects, again instructed to attend to one of two male speakers, however, now with background babble noise at different signal-to-noise ratios. Furthermore, per subject, different angular speaker positions are combined (i.e., different angular separation between the competing speakers): −90° versus +90°, +30° versus +90°, −90° versus −30°, and −5° versus +5° (see Fig. 2). More details can be found in [26]. This second dataset allows us to validate the decoding of the directional focus of attention for different angular separations and is used in Section IV-C and IV-D. Both datasets are recorded using a *C* = 64-channel BioSemi ActiveTwo system, using a sampling frequency of 8192 Hz.

**Fig. 2:**
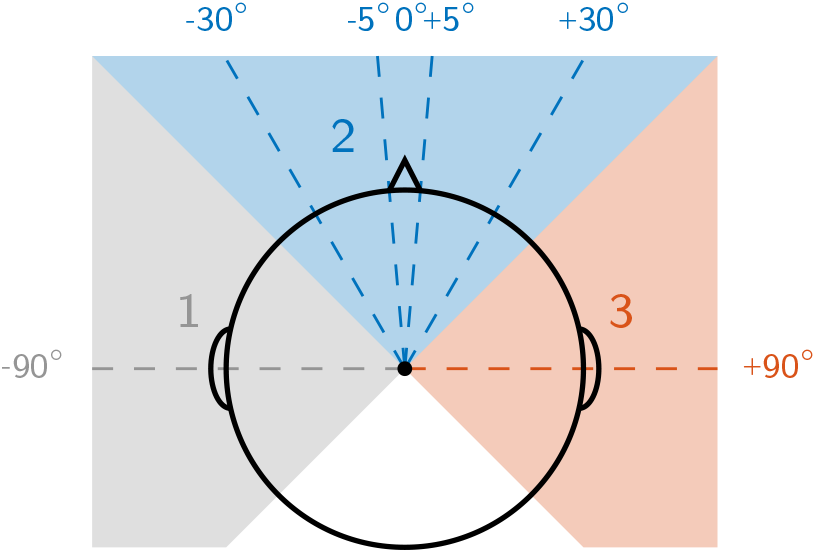
The competing speakers of Dataset II are located at different angular positions. The azimuth plane is divided into three angular domains, which are used in the multiclass problem of Section IV-D.

### B. Design choices

#### 1) EEG bandpass filtering

Before CSP filtering, a filter-bank is applied to the EEG, consisting of *B* = 14 8^th^-order Butterworth filters. The first filter corresponds to frequency band 1−4 Hz, the second to 2−6 Hz, the third to 4−8 Hz. This continues, with bands of 4 Hz, overlapping with 2 Hz, until the last band 26−30 Hz. In this way, a similar range of frequencies is covered as in [19]. The group delay is compensated for per filter using the filtfilt-function in MATLAB, resulting in a zero-phase filtering. Afterwards, the EEG data is downsampled to 64 Hz. No further preprocessing or artifact rejection is applied, as the CSP filters already implicitly suppress EEG content that is irrelevant for discrimination between both classes through a spatial filtering per frequency band.

#### 2) Covariance matrix estimation

To avoid overfitting in the estimation of the class covariance matrices in (2), the sample covariance matrices in (1) are regularized using ridge regression:

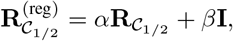

with 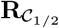 the sample covariance matrix from (1) and **I** ∈ ℝ^*C×C*^ the identity matrix. The regularization parameters *α* and *β* are not estimated using cross-validation (CV) but are analytically determined (details in [31]). This method has proven to be superior in various BCI applications and is the recommended state-of-the-art covariance matrix estimator [21].

#### 3) CSP filter design

As described in Section II-A, traditionally, the generalized eigenvalues are used to select an appropriate subset of filters, as they represent the relative output energies of each spatially filtered signal. However, these generalized eigenvalues can be influenced by outlier segments with a very high variance, which consecutively, can (negatively) affect the selection of the CSP filters. To avoid this issue, the filters are selected based on the ratio of *median* output energies (RMOE) between both classes [20], taken over all training windows with length equal to the maximal decision window length that is used in the analysis.

Furthermore, *K* = 6 CSP filters, corresponding to the 3 most discriminative filters for one and the other direction, are selected based on the cut-off point on the plot of sorted RMOEs.

### C. Performance evaluation

The FB-CSP classification pipeline is first tested per subject separately using ten-fold CV. The data per subject are therefore split into segments of 60 s (30 s for dataset II) and randomly shuffled into ten folds. This division into segments is performed in order to be able to do random shuffling over time, such that the impact of factors such as fatigue is minimized. Only in Section IV-B and the Supplementary Material, a leave-one-story+speaker-out CV (LOSSO-CV) is performed, retaining the chronological order of the segments that originate from continuously recorded trials. For each 60/30 s segment, the mean is set to zero per channel. Furthermore, each segment is normalized over all channels at once (the Frobenius norm is set to one), to assign equal weight to each segment in the training phase. During the testing phase, the normalized 60/30 s segments are split into shorter sub-segments, referred to as ‘decision windows’ (of which the length will be varied). The significance level for the accuracy is determined via the inverse binomial distribution [6].

In [8], the importance of evaluating AAD algorithms on different decision window lengths, i.e., the amount of data used to make an AAD decision, has been stressed. In typical AAD algorithms, a trade-off exists between the decision window length and accuracy. In [8], the optimal trade-off is determined by means of a criterion based on the expected time it takes to perform a stable gain switch in an attention-steered gain control system. Based on a stochastic model for the latter, and for any given decision window length, the expected time to switch the gain between speakers is minimized under the constraint of guaranteeing a pre-defined level of ‘stability’ to avoid spurious gain switches due to errors in the AAD decisions. The latter is achieved by increasing the number of gain levels, thereby increasing the gain switch duration. The optimal trade-off point between decision window length and accuracy is found as the one that leads to the shortest expected switch duration under this model, which is referred to as the *minimal expected switch duration* (MESD). The MESD is a single-number metric, facilitating the use of statistical tests to compare different AAD algorithms, as it resolves the inherent trade-off between decision window length and AAD accuracy. We refer to [8] for more details.^2^ In [8], it was found that the optimal decision window length selected within the computation of the MESD consistently shows the importance of *short* decision window lengths (*<* 10 s), allowing faster and more robust switching between speakers despite the lower AAD accuracy. To determine the accuracies on shorter decision window lengths, the left-out segments are split into shorter decision windows, on which the testing routine in Fig. 1 is applied. Note that the MESD is a theoretical metric and only provides a theoretical prediction on how an optimized AAD-based gain control algorithm would track attention switches [8]. Here, we do not experiment with data containing actual attention switches.

The hyperparameters of the LDA classifier are optimized on the CSP output energies of the training set using five-fold CV.

## IV. Results and discussion

### A. Comparison with stimulus reconstruction approach

The FB-CSP classification method is compared to the current state-of-the-art AAD method, which adopts the stimulus reconstruction approach, on Dataset I. Here, canonical correlation analysis (CCA) is used, which is considered to be one of the best decoding methods to date, outperforming other backward and forward models [2]. In CCA, a jointly forward (i.e., mapping the stimulus envelope to the EEG) and backward (i.e., mapping the EEG to the stimulus envelope) model is trained and applied to new data [24]. The attended speaker is identified by classifying the difference between the canonical correlation coefficients of the competing speakers using an LDA classifier. A forward lag of 1.25 s is used on the speech envelopes and a backward lag of 250 ms is used on the EEG as in [2], [24]. The CCA method is tested using the same ten-fold CV procedure as for the FB-CSP method. The number of correlation coefficients used in the LDA classification is determined by an inner ten-fold CV loop. No a priori principal component analysis or change of filter basis as in [24] is used. The EEG and speech envelopes, which are extracted using a power-law operation with exponent 0.6 after subband filtering [7], are filtered between 1 and 9 Hz (thus mainly without *α/β*-activity, which was determined to be optimal for linear stimulus reconstruction [6], [7], [11], [26], [29]) and downsampled to 20 Hz. Note that this method employs an inherently different strategy for AAD than FB-CSP, by (in a way) reconstructing the attended speech envelope, rather than decoding the directional focus of attention.

In Fig. 3a, it is observed that this stimulus reconstruction approach is characterized by a degrading accuracy for shorter decision window lengths, while the accuracy of the FB-CSP method barely decreases. It thus clearly outperforms the stimulus reconstruction approach for short decision window lengths. This is one of the most important properties of this new strategy for AAD to decode the directional focus with FB-CSP rather than reconstructing the stimulus. This effect was also seen in [19], where the directional focus is decoded based on a CNN. While the stimulus reconstruction approach tries to determine the attended speaker by reconstructing the attended speech envelope, the FB-CSP method only needs to *discriminate* between two angular directions, which is an inherently easier filter design strategy. Furthermore, in the former, correlation is used as a feature, of which the estimation is inaccurate when computed on short decision windows, in particular because the correlation coefficients observed in stimulus reconstruction are very small, making their estimation susceptible to noise. Lastly, as mentioned before, the FB-CSP method is mostly based on an instantaneous spatial feature (brain lateralization) rather than a temporal feature.

**Fig. 3:**
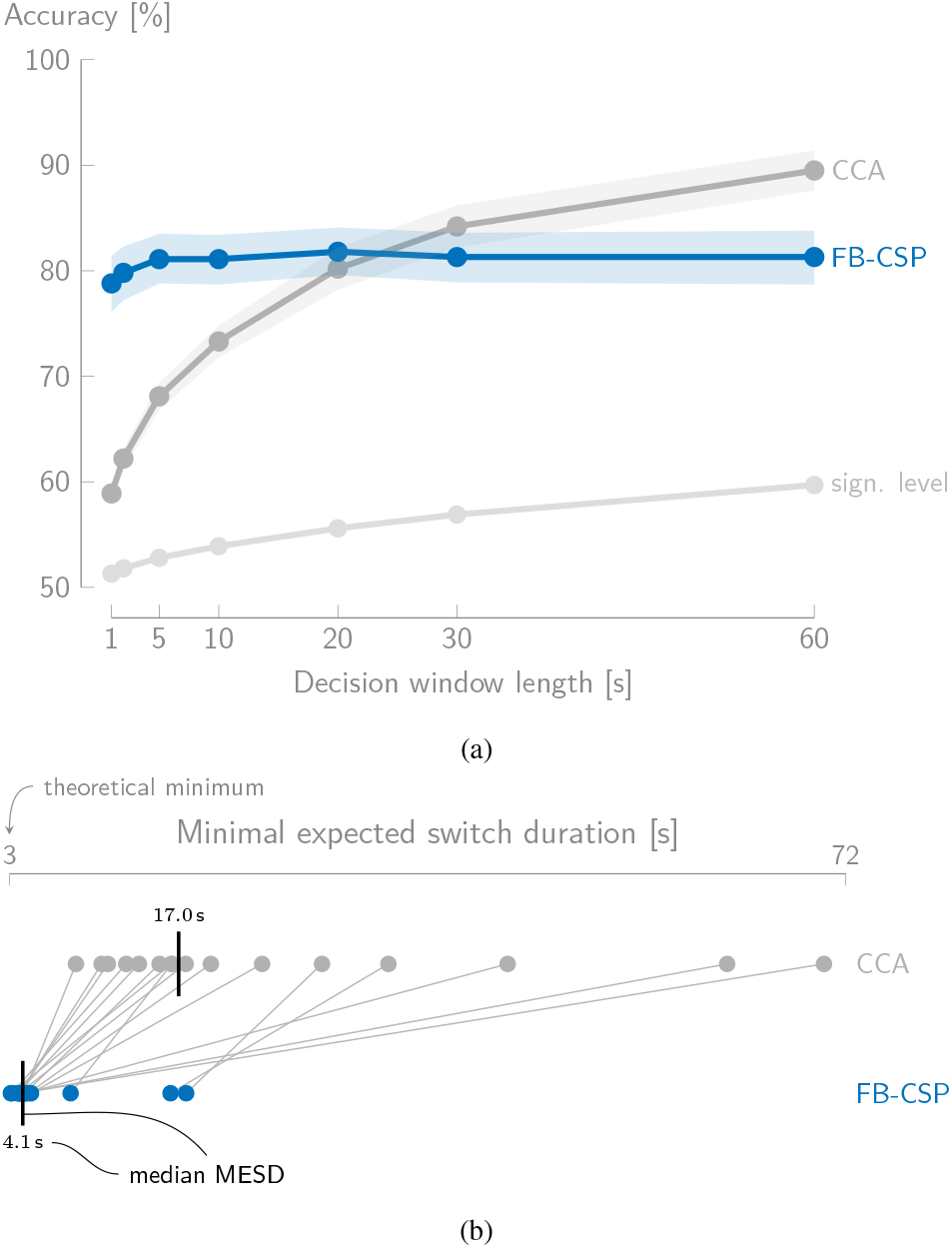
(a) The accuracy (mean ± standard error of the mean across subjects; Dataset I) of the FB-CSP classification method barely decreases for shorter decision window lengths and outperforms the stimulus reconstruction approach (CCA) for short decision window lengths. Note that the significance level (‘sign. level’) decreases for shorter decision window lengths due the higher number of test windows. (b) The median MESD (black vertical line) is significantly lower (better) for the FB-CSP method than for the stimulus reconstruction approach (CCA). Each dot represents one subject, the lines connect the same subjects across methods.

Note that the accuracy of the FB-CSP method exhibits a higher inter-subject variability than the stimulus reconstruction method. We do not consider this as a major disadvantage of the FB-CSP method, as, for example, on 1 s decision windows, performance is still better for *all* subjects compared to CCA. On average, there is a 20% gap in accuracy for 1 s decision windows.

For long decision window lengths, however, CCA outperforms the FB-CSP method. To resolve this trade-off and to (statistically) determine which method performs better in a context of neuro-steered gain control for hearing devices, we use the MESD metric [8], a relevant criterion for AAD that optimizes the speed-accuracy trade-off (Section III-C) and thus resolves the inconclusiveness based on the performance curve. Fig. 3b shows the MESDs per subject, for both algorithms. It is clear that FB-CSP (median MESD 4.1 s) results in much faster switching than CCA (median MESD 17.0 s). A Wilcoxon signed-rank test confirms that there indeed is a significant difference between the MESD for the FB-CSP method versus CCA (*W* = 0, *n* = 16, *p* < 0.001). The sustained performance for short decision window lengths thus results in a superior performance (for all subjects) of the FB-CSP method over CCA. We note that the MESD is by definition longer than the decision window length (see Section III-C and [8]). In particular, the theoretical lower limit for the MESD is 3 s when using a minimal decision window length of 1 s^3^ [8].

### B. Comparison with convolutional neural network approach

In [19], a convolutional neural network (CNN) is used to perform the same task, i.e., decoding the directional focus of attention. This CNN approach has been validated on the same dataset (Dataset I), but with a different testing procedure to avoid overfitting on speakers, i.e., LOSSO-CV instead of random CV. To provide an honest and transparent comparison of our FB-CSP method with this CNN method, we have cross-validated the performance of the FB-CSP method in the same way as in [19], at the cost of less training and testing data. While data of other subjects are included in the training of the CNN method as a regularization technique [19], this is not done for the FB-CSP method. The EEG data are filtered between 1 − 32 Hz, as proposed in [19] and equivalent to the FB-CSP method.

Given the performances in Fig. 4, we, first of all, want to stress that the results of the FB-CSP method for a LOSSO-CV are very similar to using a random CV (Fig. 3). This confirms that, as opposed to the CNN method, our FB-CSP method does not overfit on speakers or stories, which could occur when using random CV. For the CNN method, the results were significantly better when not leaving out the speaker and/or story in the training set, which could be a sign of overfitting [19]. Furthermore, our FB-CSP method does not perform worse than the CNN method, as a Wilcoxon signed-rank test (*W* = 56, *n* = 15, *p* = 0.85, one outlier subject removed) shows no significant difference based on the MESD (Fig. 4b).

**Fig. 4:**
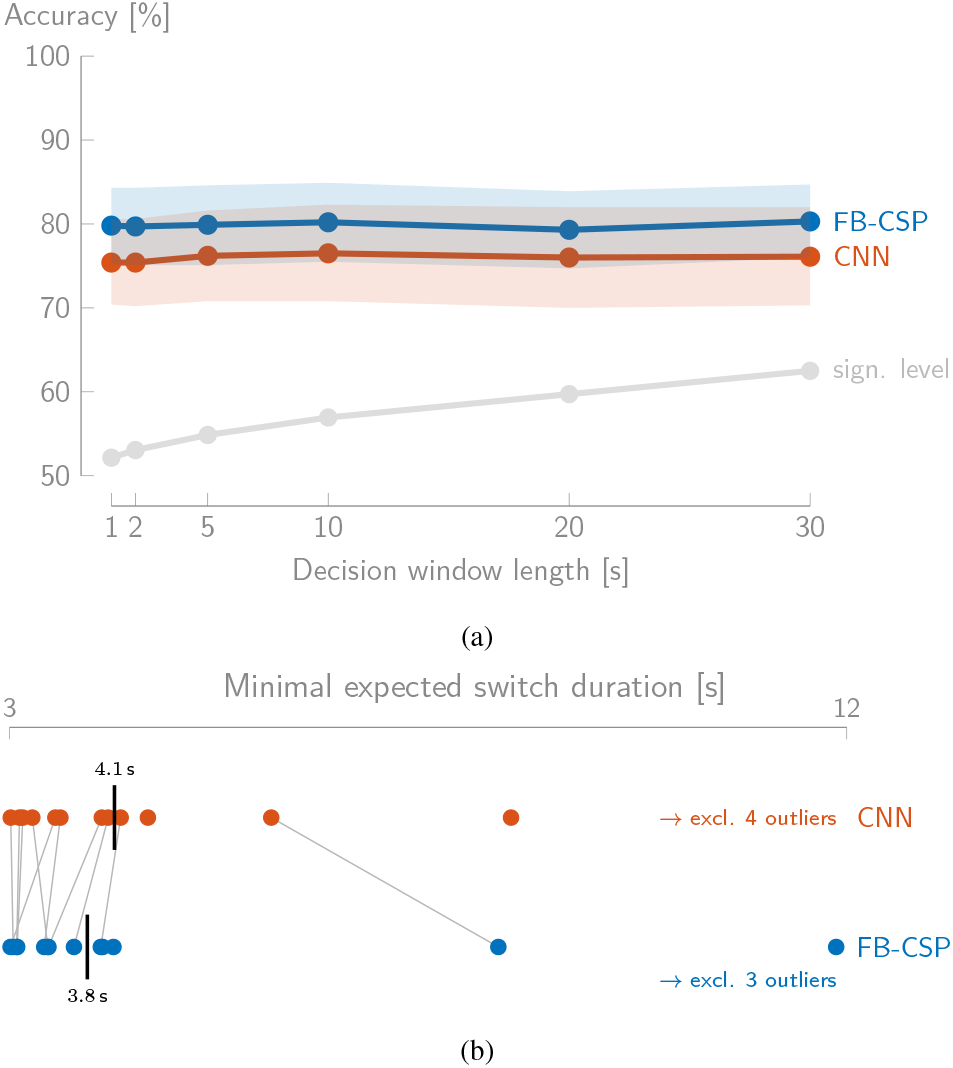
(a) The FB-CSP method outperforms the CNN method with on average ≈ 4% in accuracy (mean ± standard error of the mean; Dataset I). (b) The median MESD is lower for the FB-CSP method than for the CNN method. Note that when the MESD of a subject is not connected to the corresponding MESD, it corresponds to an outlier value of the other method.

To conclude, we have identified the following advantages of the FB-CSP method over the CNN method:

- The FB-CSP method does not perform worse than the CNN method, it even tends to outperform it.
- The FB-CSP method shows less inter-subject variability and is more stable (see the standard error of the mean in Fig. 4a and the spread in Fig. 4b).
- The FB-CSP method requires less training for a better performance. The CNN method uses training data of all (other) subjects, including the test subject, to avoid overfitting.
- The FB-CSP method has a lower computational complexity, which is paramount to be applicable in mobile and wireless hearing devices.

### C. Binary FB-CSP classification at various speaker positions

Whereas in the previous experiments, the competing speakers are located at −90°/+90°, this section treats binary AAD classification at the various speaker positions that are present in Dataset II. Fig. 5 shows the performance of the FB-CSP classification method. For each pair of competing speaker positions, all babble noise conditions are pooled and the FB-CSP classification method is applied. Each pair of positions is thus treated separately, with independently trained CSP filters and LDA classifiers.

**Fig. 5:**
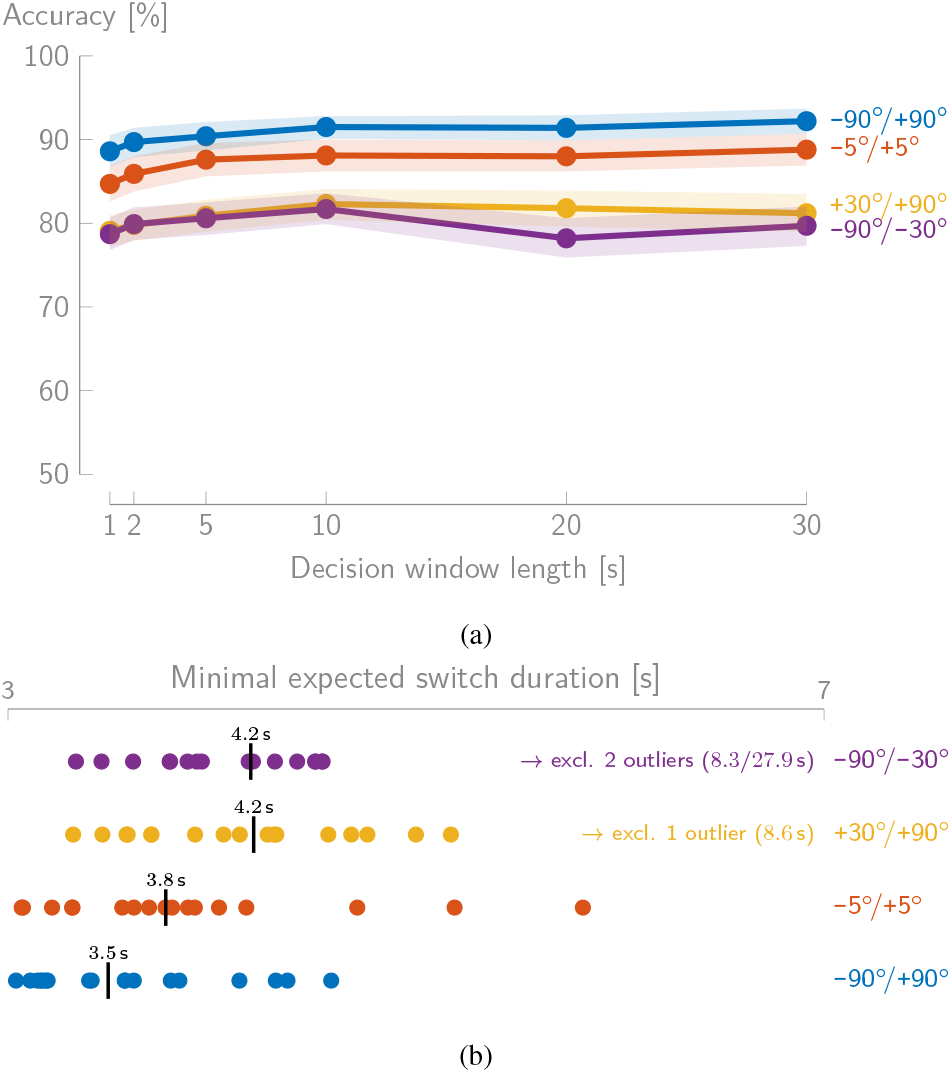
(a) The FB-CSP classification method performs well for all speaker separation angles (Dataset II). Again, the accuracy (mean ± standard error of the mean) of the FB-CSP classification method barely decreases for shorter decision window lengths. (b) The median MESD is lower (better) for the −90°/+90° than for the other scenarios. Decoding the directional focus of attention for speakers that are positioned at the same side of the head is harder than when they are symmetrically positioned on different sides of the head.

First of all, the results in Fig. 5a confirm and reproduce the previous results from Fig. 3a. The accuracy for the −90°/+90° condition is on average even 10% higher and there is a smaller inter-subject variability (standard deviation is on average over all decision window lengths ≈ 7%, while this was ≈ 10% in Dataset I). A possible explanation for this difference in performance is that due to the presence of background noise, the spatial cues become more important; or that the subject has to focus harder, thereby generating stronger neural responses. A similar advantageous effect of the presence of background noise was observed in [26].

Furthermore, these results allow analyzing the effect of the angular speaker separation on the decoding performance. The main result from these performances is that decoding the directional focus of attention from the EEG using the FB-CSP classification method still works for various angular scenarios, and even when the speakers are positioned very closely together (−5°/+5°) or are positioned at the same side of the head (+30°/+90° and −90°/−30°). The MESDs in Fig. 5b confirm these findings.

As can be expected, the decoding for the −90°/+90° scenario is easier than in the other scenarios. Although the decoding will fail when speakers are co-located at the same spatial position, the FB-CSP method still succeeds in reliably discriminating between very closely positioned speakers at −5°/+5°. Furthermore, as the results for −5°/+5° are still better than when the competing speakers are positioned at the same side of the head, it seems that when speakers are located at different sides of the head, this provides a substantial advantage in decoding the directional focus of attention. However, even when speakers are located at the same side of the head, the method finds sufficient spatio-temporal discriminative patterns to differentiate between speaker locations.

As an important consequence of these results, the FB-CSP method can be used as a basic building block for a new AAD strategy in which, for example, the whole plane along the azimuth direction is split into angular domains. Depending on the multiclass coding strategy, several FB-CSP filters are then combined to locate the attended speaker in the plane and to steer a beamformer into the correct direction. This AAD strategy is tested in the following section.

### D. Multi-condition and -class FB-CSP classification

Using Dataset II, we can verify whether a multi-condition or -class strategy is feasible. In the first experiment, all data are pooled and the FB-CSP classifier tries to determine whether the *right- or left-most* speaker is attended. In the second experiment, all angles are divided into three angular domains (left/frontal/right) as depicted in Fig. 2.

#### 1) Classifying the right/left-most speaker as attended speaker

Instead of training an FB-CSP for each angular condition separately, all conditions can be pooled and the FB-CSP classifier can be trained to determine whether the user is listening to the right-most or left-most speaker (in a two-speaker scenario), *independent* of where these speakers are positioned in the plane. As a consequence, a speaker positioned at −30° (which is located at the left side of the head) can be the right-most attended speaker, relative to −90°, while +30° (which is located at the right of −30°) can be the left-most attended speaker, relative to +90°. This angular condition-independent FB-CSP classifier could then be used generically to steer a beamformer or to select the attended speaker, provided the angular positions of the competing speakers are known or can be detected from a hearing device’s microphone array. In order to test this, all the data of Dataset II are pooled and randomly divided into ten folds. Note that a limitation of this experiment is that the different speaker positions only appear in fixed pairs and that not every position is combined with all other positions.

Fig. 6 shows that the accuracy when classifying attention to the left/right-most speaker is still high (77.7% on average over all decision window lengths), although lower than when classifying each condition separately (Fig. 5a). This confirms that this strategy is viable.

**Fig. 6:**
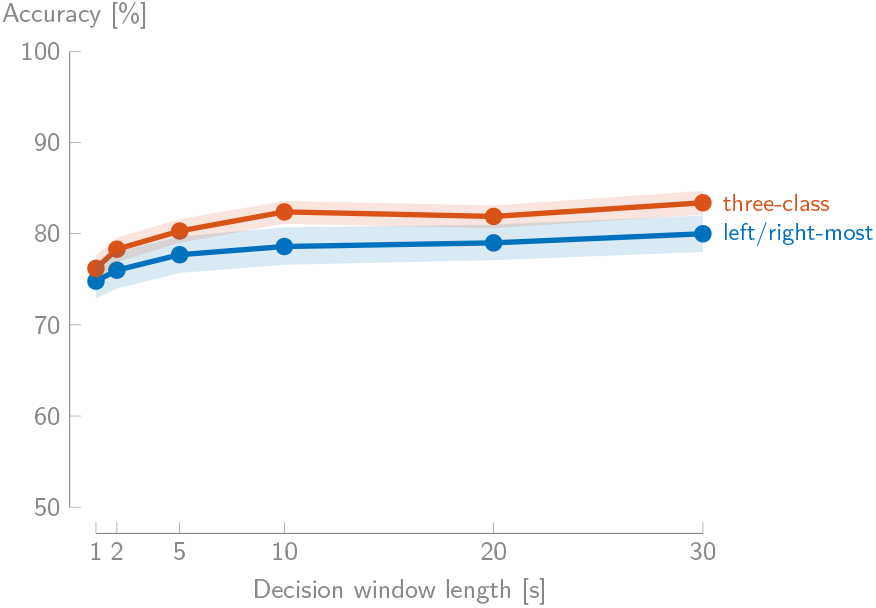
These peformance curves show that even when pooling all conditions and only classifying the attention to the left/right-most speaker, as well as dividing the upper half-plane in three angular domains as in Fig. 2, the accuracy (mean *±* standard error of the mean; Dataset II) is still high.

When investigating the MESDs per angular condition (still when classified all together), it is clear that there are two groups (Table I): the first group contains the conditions where the competing speakers are located along different sides of the head and show only a small increase in MESD compared to when they are classified separately (compare with Fig. 5b), while there is a larger increase in MESD when the competing speakers are positioned at the same side of the head. Furthermore, the first group shows a lower MESD than the latter one.

**Tab I:**
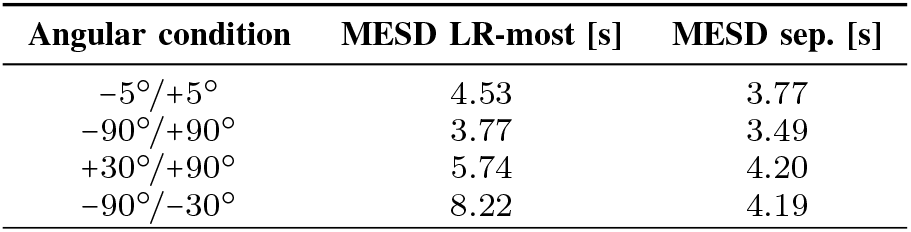
The median MESD is lower when the speakers are located on different sides of the listener. Furthermore, the MESDs are higher compared to the case where each condition is classified separately (MESD sep.; see Fig. 5b).

#### 2) Classifying between left/frontal/right direction

The intuitive multiclass extension of the binary classification of only two angular conditions is to classify multiple speaker positions at the same time, i.e., determining the directional focus of attention among several possibilities. A possible strategy could be to divide the azimuth plane into different angular domains, which are classified together. In this way, a beamformer could be steered towards the correct angular domain (without having to also estimate the direction of arrival of each speaker separately from the microphone recordings). The higher the spatial resolution of the multiclass strategy, the lower the chance that multiple speakers are present in the same angular domain (in case of multiple competing speakers), but the higher the misclassification error. In case multiple speakers are detected within each angular domain, more angle-specific classifiers or the aforementioned strategy of classifying the left/right-most classifier (Section IV-D1) could be used as a complementary approach.

To test the feasibility of this strategy, we divide the azimuth plane into three classes based on speaker position as in Fig. 2. The segments in Dataset II are divided into these classes accordingly. Note that the same limitation as before (limited speaker pairs) holds here and that there are no other positions present than ±90° in domains 1 and 3. A one-versus-all coding scheme is used, which means that there are three binary classifiers trained, which each classify one angular domain versus the other two domains combined.

Fig. 6 shows the performance curve for this three-class problem. The accuracies are very high and show low subject-to-subject variability (standard deviation ≈ 5.6% over all decision window lengths). Note that the accuracy decreases faster for shorter decision window lengths than usual. This effect is to a lesser extent also present in the binary case and is amplified here because of the multiclass nature of this problem. However, the decrease is still very limited and results in short switch durations (median MESD of 4.32 s over all angular domains, 4.58 s for switching to domain 1, 4.01 s for switching to domain 2, and 5.13 s for switching to domain 3).

### E. Channel selection

For the FB-CSP method to be applicable in the context of neuro-steered hearing devices, which is an inherently mobile application, we test the method with a reduced set of EEG channels. However, we do not adopt a traditional data-driven feature/channel selection method but take an application-based point of view. The five electrodes closest to each ear are selected from the 64-channel BioSemi system (see the blue channels on Fig. 7). This can be viewed as a representative selection that mimics current behind-the-ear EEG approaches such as the cEEGrid array [32], which has also been used for AAD [33]. However, it is noted that our analysis is not fully representative of an actual cEEGrid setup due to different recording equipment and different electrode positions. We mainly want to verify whether decoding the directional focus of attention while dominantly measuring from the electrodes on the temporal lobe, is possible.

**Fig. 7:**
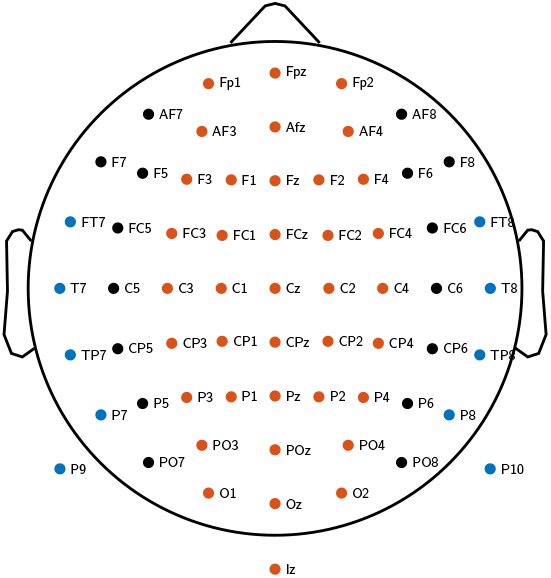
The five electrodes of the 64-channel BioSemi system closest to the ear are selected for the channel selection in Section IV-E (blue). In Section IV-H, 38 central electrodes are chosen (orange).

To eliminate the dependence on an ‘external’ or joint reference electrode, the selected EEG channels are re-referenced using a common average reference for each ear separately. By averaging and re-referencing per each ear separately, the two sets of ear channels are galvanically isolated, i.e., emulating two standalone EEG sensor devices which do not have to be connected with a wire. Furthermore, common average referencing is used to eliminate the need for selecting a particular reference electrode. Per ear, one random (as CSP filtering is invariant to the removed channel) re-referenced EEG channel is removed to avoid rank-deficiency in the EEG covariance matrices, effectively leading to 4 channels per ear (*C* = 8). After the removal of the other channels and the referencing, the complete FB-CSP pipeline (Fig. 1) is retrained and evaluated using the reduced set of EEG channels.

Fig. 8a shows that the decrease in accuracy on Dataset I (binary classification) when selecting the ear channels is limited to ≈ 5.6% on average. Furthermore, the median MESD increases from 4.10 s (64 channels) to 4.74 s, which is statistically significant (*W* = 0, *n* = 16, *p <* 0.001), but is still limited. Lastly, from Fig. 8b, it can be seen that there is only a limited increase in variability over subjects.

**Fig. 8:**
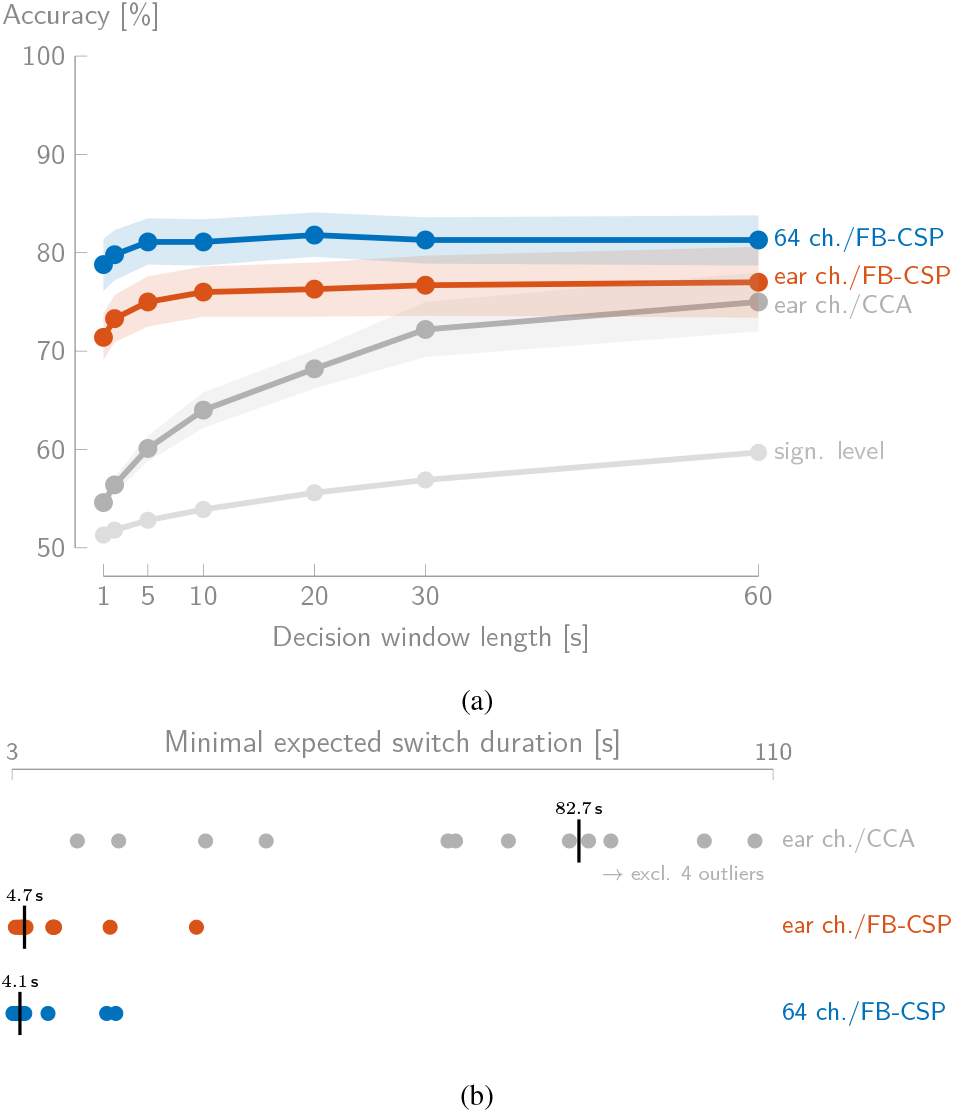
(a) The mean accuracy (± standard error of the mean) when using only ten electrodes close to the ear on Dataset I (binary classification) decreases relatively little compared to the full 64-channel setup. (b) There is a limited increase in median MESD when selecting ten electrodes for the FB-CSP method, while the CCA method greatly suffers from the channel reduction (compared to Fig. 3b).

Furthermore, also the performance of CCA is shown, using the same reduced set of channels and corresponding referencing method. The accuracy decreases on average with ≈ 10% over all decision window lengths (Fig. 3a) and does not outperform the FB-CSP method anymore on long decision window lengths. The median MESD drastically increases as well (Fig. 8b). The stimulus reconstruction approach thus suffers more from the channel reduction and is completely outperformed by the FB-CSP method, which is another advantage of the newly proposed method.

We conclude that decoding the directional focus of attention with the FB-CSP method using a reduced set of channels close to the ear could be possible, but that there is more research required to further validate this approach.

### F. Performance on very short decision window lengths (< 1 s)

Fig. 9 shows the performance of the FB-CSP method on Dataset I (binary classification) for 64 channels and the channels close to the ear (see Section IV-E) for decision window lengths below 1 s. Below 1 s, the accuracy further degrades, with a limited loss of ≈ 5.5% accuracy on 31.25 ms decision windows and ≈ 8.5% on 15.63 ms decision windows compared to 1 s decision windows. As a result, for both setups, there still is an acceptable performance when taking quasi-instantaneous decisions, resulting in a median MESD of 76.5 ms (64 channels) and 195.0 ms (ear channels) over all subjects. Note that caution is needed when interpreting these MESD values, as on such short decision window lengths, the independence assumption of the Markov model underlying the MESD metric is gravely violated due to the significant autocorrelation values of EEG signals below 1 s lags. The actual time to achieve a sufficiently stable switch may be slightly higher than the one predicted by the model behind the MESD metric.

**Fig. 9:**
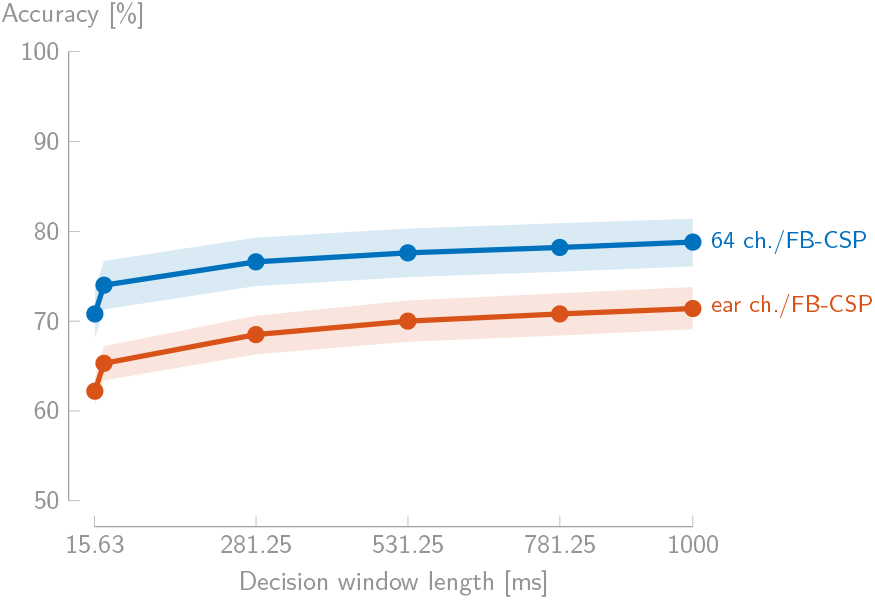
The performance curves (mean ± standard error of the mean) of the FB-CSP method degrade below 1 s decision window lengths, while demonstrating acceptable performance even for quasi-instantaneous decisions (Dataset I, binary classification).

While it may seem surprising that the method can still decode the direction of attention quasi-instantaneously (*<* 32 ms) with an accuracy that is better than chance, we note that CSP only exploits spatial information (differences between channels) rather than temporal information. Integrating over a longer time window only helps to achieve a better estimate of the log-energies that are fed to LDA, which is the reason behind the slight increase in performance for longer decision windows (compared to instantaneous log-energy estimates). In the case of CSP, the length of the decision window is less important than in stimulus reconstruction approaches, where temporal modulations in the speech envelopes are exploited and where the decision window length directly determines how much of this information is available for discrimination between both speakers. Furthermore, in the case of FB-CSP, the estimation errors on the log-energies (due to quasi-instantaneous estimation) can be further compensated by the LDA classifier by exploiting redundancy in the different filter bands and CSP components to make a reliable decision. Lastly, although the FB-CSP method makes a decision based on a few samples, because of the filterbank on the EEG, these samples are also the result of a weighted integration of previous samples. This means that effectively more samples than the number of samples in the decision window are used.

### G. CSP classification on an unseen subject

In the preceding experiments, the FB-CSP filters and LDA classifiers are trained subject-specifically. Here, we test the viability of the subject-independent approach of Section II-D, to improve the practical applicability of this method in neuro-steered hearing devices. The same (FB-)CSP classification pipeline (Fig. 1) and design choices (Section III-B) as before are used, but now tested on Dataset I (binary classification) in a *leave-one-subject-out* manner. Per test subject, the (FB-)CSP filters and LDA classifier are trained on the 15 other subjects. Without using any of the adaptions from Section II-D, the subject-independent FB-CSP method (SI-FB-CSP) exhibits a large drop in performance in comparison with the subject-specific FB-CSP method (SS-FB-CSP) (see Fig. 10a).

**Fig. 10:**
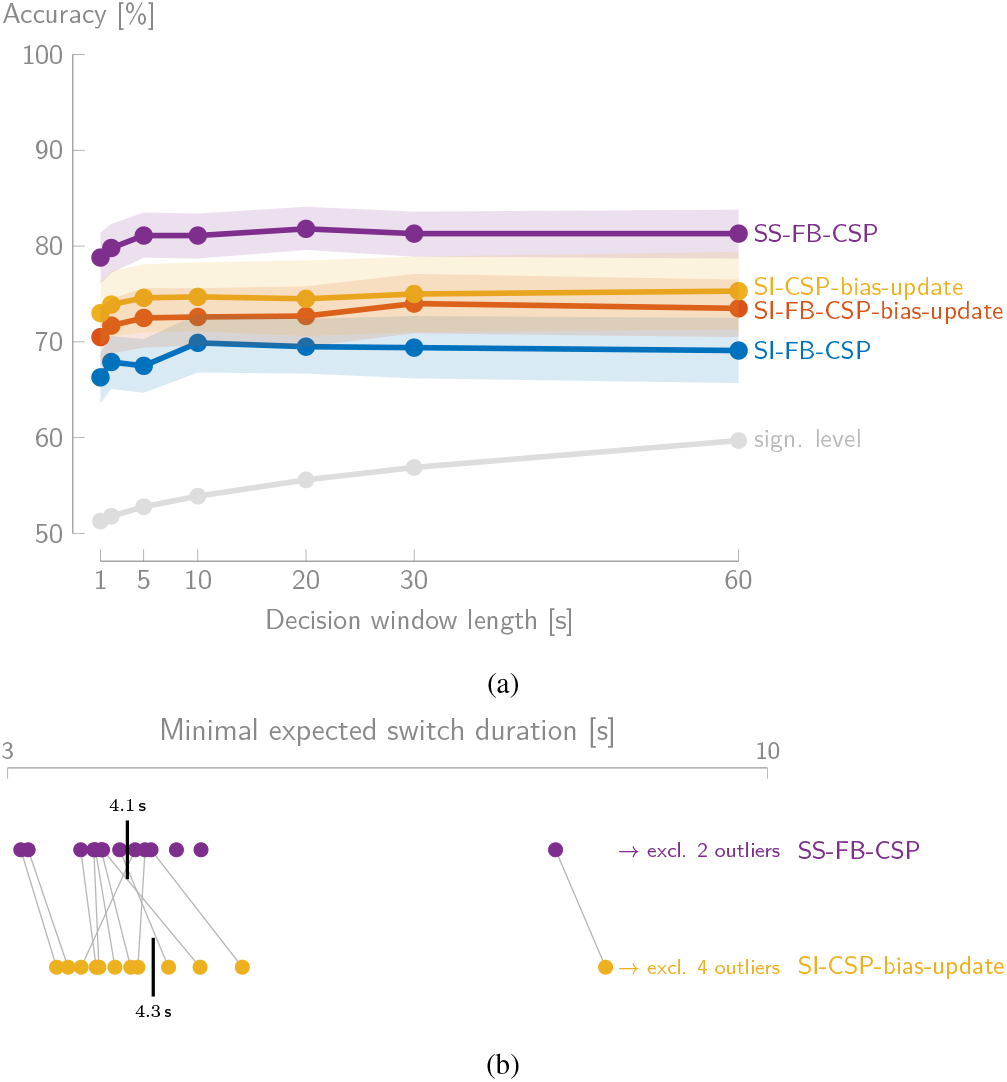
(a) Using a bias update (SI-FB-CSP-bias-update) and only one frequency band (SI-CSP-bias-update) in the subject-independent CSP classification method on Dataset I (binary classification) results in a substantial increase of performance over the baseline (SI-FB-CSP) (mean ± standard error of the mean). (b) The median MESD of the subject-independent CSP classifier (SI-CSP-bias-update) is very close to the one of the subject-specific FB-CSP classifier (SS-FB-CSP). There is, however, a larger spread, with more negatively (higher) outlying MESD values. Note that when the MESD of a subject is not connected to the corresponding MESD, it corresponds to an outlier value of the other method.

Updating the bias as in Section II-D results in a substantial increase of performance of ≈ 4% (SI-FB-CSP-bias-update). The second adaptation reduces the FB-CSP method to a CSP method by using a single frequency band (the *β*-band: 12-30 Hz, *B* = 1), which was experimentally determined (see Section IV-H). Using this CSP method in combination with a bias update of the LDA classifier results in another increase of accuracy (Fig. 10a; SI-CSP-bias-update).

The best subject-independent CSP classifier, with a bias update and only one frequency band (SI-CSP-bias-update), is compared with the subject-specific FB-CSP classifier (SS-FB-CSP) in Fig. 10a and Fig. 10b. Note that using a single frequency band for the subject-specific method (SS-CSP) results here in a ≈ 2% decrease in accuracy over all decision window lengths. From the MESD, we can see that the subject-independent method quite nicely approximates the performance of the subject-specific method. For two subjects, the subject-independent method even performs better than the subject-specific FB-CSP method. However, there still is a significant difference (Wilcoxon signed-rank test: *W* = 127, *n* = 16, *p* = 0.0023). Furthermore, from Fig. 10b, it can be seen that the subject-independent method has a larger spread, with more negative outlier values.

We conclude that the subject-independent CSP classification on average approximates the performance of a subject-specific FB-CSP classifier in terms of MESD, but that there is no guarantee that it will work on every subject. This slightly worse performance is, however, traded for practical applicability, as no a priori calibration session per user is required.

### H. Decoding mechanisms

Given that it is possible to decode the directional focus of attention with CSPs, it is relevant to get a handle on what drives the decoding. To investigate which frequency bands are most important, the subject-independent FB-CSP pipeline is trained on all subjects with *B* = 4 filter bands, corresponding to the main EEG frequency bands (1 — 4 Hz (*δ*), 4 — 8 Hz (*θ*), 8 — 12 Hz (*α*), and 12 — 30 Hz (*β*)). The mean leave-one-subject-out accuracy over all subjects using a 60 s decision window length is 79.7%^4^. To assess the importance of each band, the *K* = 6 energies related to each band are left out (while keeping all others), leading to a decrease in accuracy to 79.0% for the *δ*-band, 79.3% for *θ*-band, 79.0% for the *α*-band, and 73.2% for the *β*-band. This indicates that the *β*-band is the most important band, motivating the choice of this band in Section IV-G. Similar conclusions have been drawn in [19], [34]. Furthermore, the performance does not degrade over time when the attention is sustained (see Supplementary Material), which has been reported in the context of *α*-power lateralization [17].

Fig. 11 shows the spatial activations of the *β*-band CSP filters. These topographic maps show activations mainly above the fronto-temporal cortex, consistent with the *β*-band activity found in [19], [34]. However, caution is needed when interpreting these spatial maps: the CSP filters implement a so-called ‘backward’ decoding model, which could implicitly also perform suppression of non-related EEG activity and artifacts, and can thus result in misleading interpretations [35]. To make the spatial maps as interpretable as possible, eye (blink) artifacts have been removed with ICA and muscle artifacts have been removed with CCA [36], making it impossible for the CSP filters to reconstruct and exploit them. Note that because of the artifact removal, the spatial filters shown in Fig. 11 do not correspond to the ones applied in the experiments. We merely try to highlight the neural underpinnings of the spatial filters by removing artifact-related activity before computing the CSP filters. As such, the topographic plots are not affected by the artifact removal mechanism that would normally be present implicitly in the CSP filters themselves (i.e., if such an implicit artifact removal would help in maximizing the discrimination between the classes). We reiterate that (linear) artifact removal is unnecessary in the experiments, as the CSP filters can deal with artifacts. Furthermore, the subject-specific performance on 60 s windows, using only the *β*-band, with and without mentioned artifact removal is very similar (77.2% vs 79.0% respectively), which indicates that the artifacts are not harming nor driving the CSP-based decoder.

**Fig. 11:**
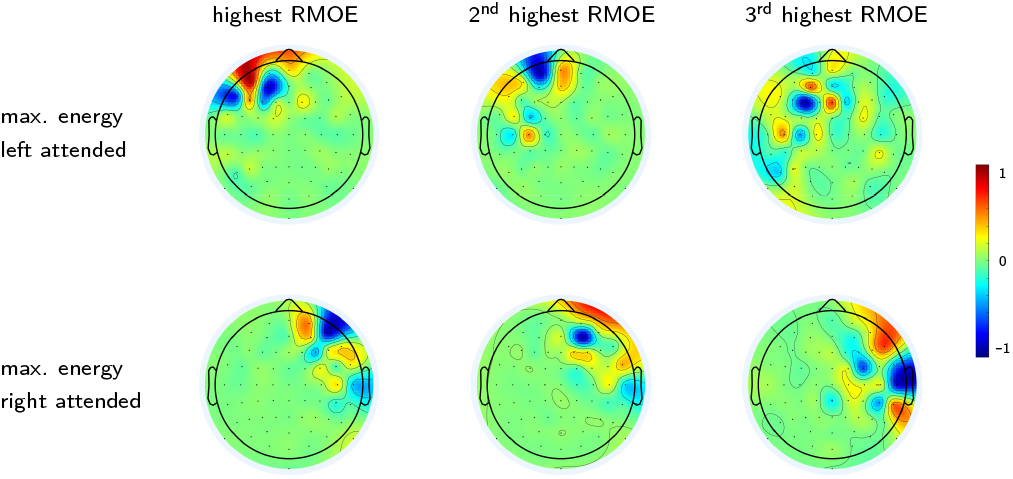
The topographic plots of the six spatial *β*-band CSP filters, computed on all data of all subjects of Dataset I, show mainly fronto-temporal activity. The filters of the first row maximize the output energy when left is attended, while those in the second row maximize the output energy when right is attended. The columns correspond to different RMOEs.

Whether the CSPs exploit neural information or some correlated artifact signal (eye artifacts, EMG, …) is impossible to determine. Intuitively, two specific types of artifact signals could be potentially exploited by the CSPs: eye artifacts (e.g., lateral movements) and EMG activity (especially subtle directive ear movements [37]). However, there are several indications that the CSP filters do *not* exploit these effects and indeed focus mainly on neural activity.

It is very unlikely that the CSPs exploit eye artifacts, as they are mostly contained in the *δ*- and *θ*-band, whereas the CSP filters focus on *β*-band activity. Secondly, the explicit removal of the eye blinks using ICA does not affect the performance (80.0% on 60 s decision windows). Furthermore, the decoding also works well when the competing speakers are located at the same side of the head (Section IV-C) and even when the subjects are asked to fixate on a cross (tested on the dataset of [38], results not shown).

As ear movements spectrally (*β*- and *γ*-band) overlap with the information used by the CPs, it is more difficult to exclude the exploitation of subtle ear movements [37]. There are, however, two counterindications. Firstly, the approach also works for speakers located at the same side of the head (Section IV-C). Secondly, when using only the 38 most central electrodes (out of the 64 channels) furthest away from the ears (see the orange electrodes in Fig. 7) and the *β*-band activity, we still obtain a subject-specific accuracy (on Dataset I) of 74.1% on 60 s windows. This at least shows that the decoding still works when only using the central channels, and thus most probably while not being able to pick up ear muscle activity. Furthermore, the topographic plots in Fig. 11 show that the CSP filters also exploit channels that are rather far away from the ears, even when the channels close to the ears are included in the data-driven design.

## V. Conclusion

We have shown that a (filterbank) common spatial patterns classification method is capable of decoding the directional focus of attention, solely based on the EEG. An inherent limitation of this approach is that it requires the competing speakers to be spatially separated. Furthermore, this spatial separation needs to be perceived by the user, which is more difficult for certain hearing impaired populations.

The proposed method has shown to not only outperform the classical stimulus reconstruction approach for auditory attention decoding in a two-speaker situation but does also not perform worse than a computationally more complex convolutional neural network approach that performs the same task [19]. It achieves practically viable MESDs below 4 s, which has not been achieved by any other AAD method so far [2]. Furthermore, the proposed method has several important advantages, which are important for practical use in neuro-steered hearing device applications:

1. the FB-CSP method does not require clean speech envelopes (in contrast to the traditional stimulus reconstruction approach), such that the extra (error-prone) speech separation step for AAD can be avoided,
2. the performance barely decreases for short decision window lengths and still achieves acceptable performance for quasi-instantaneous decisions, potentially resulting in very fast and robust switching between speakers,
3. the method still works using a limited set of EEG channels above the ears,
4. the method is capable of discriminating between different angular speaker positions,
5. the method can be employed within a multi-condition or multiclass strategy to handle multiple speaker positions at the same time,
6. the method can, provided minor updates, be used in a subject-independent way, trading a minimum of performance for practical applicability.

We believe that these assets make the FB-CSP method an excellent candidate and a major step forward towards practical neuro-steered hearing devices.

## VI. Acknowledgments

The authors would like to thank Simon Van Eyndhoven for providing the implementation of the CSP filter design and the shrinkage covariance estimator.

## Supplementary material

In the supplementary material, related to the paper *Fast EEG-based decoding of the directional focus of auditory attention using common spatial patterns*, we investigate the AAD accuracy as a function of time during sustained attention.

### A. Decoding the directional focus during sustained attention

To investigate the AAD accuracy as a function of time during *sustained attention*, we use the leave-one-story+speaker-out cross-validation of Section IV-B in the paper on Dataset I, allowing to leave out full continuous recordings. It is important to verify whether decoding the directional focus of attention is possible during the *full* duration of a continuous recording, while the subject sustains its attention towards a particular speaker/direction. If the AAD accuracy degrades over time, this means the FB-CSP method only exploits brain lateralization patterns when the subject initially focuses its attention, which has been reported in the context of *α*-power lateralization [1].

Fig. 1 shows the averaged performance over continuous trials and subjects as a function of time. As Dataset I contains 6-minute continuous recordings (here referred to as trials) of EEG with sustained attention, the AAD accuracy is shown per 1 s sliding decision window (no overlap) over these trials. The mean accuracy, over all decisions, 6-minute trials, and subjects, is equal to 80.0% and is the same as the accuracy on the 1 s-point in Fig. 4a in the paper. Furthermore, there is no apparent decrease in performance over time, on the contrary, the accuracy seems to slightly increase in the first minute, whereafter the accuracy remains constant. This confirms that the FB-CSP method is capable of decoding the directional focus of attention when the attention is sustained, furthermore, with a similar accuracy as when using random cross-validation (see Fig. 3a).

**Fig. 1:**
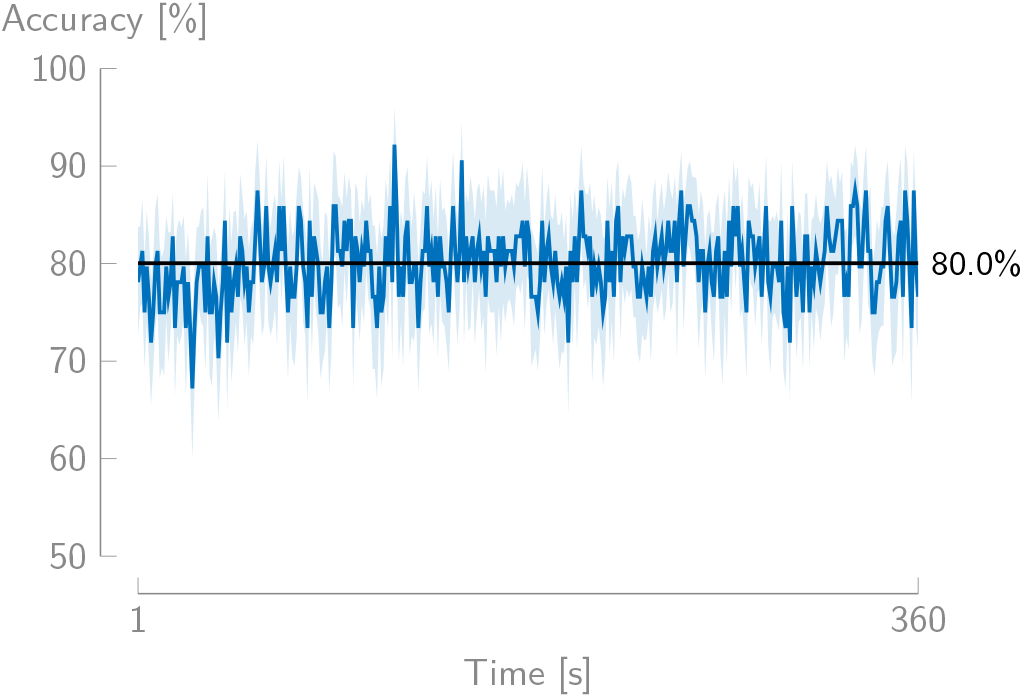
The performance (mean accuracy ± standard error of the mean over subjects and different 6-minute trials; Dataset I) does not degrade as a function of time when the attention is sustained. A sliding window of 1 s is used.

## Footnotes

1 The code for the subject-specific experiments on this dataset are available at https://github.com/exporl/spatial-focus-of-attention-csp.

2 A toolbox to compute this metric is available at https://github.com/exporl/mesd-toolbox.

3 The theoretical lower limit of the MESD is equal to 3 the shortest decision window length that is tested with, as for 100% accuracy, three steps must be taken in the Markov chain [8].

4 As the data of the subject under test is used in the CSP training (but not in the LDA training), this accuracy is slightly higher than in Fig. 10.

